# Involvement of superior colliculus in complex figure detection of mice

**DOI:** 10.1101/2022.09.25.509365

**Authors:** J. Leonie Cazemier, T. K. Loan Tran, Ann T. Y. Hsu, Medina Husić, Lisa Kirchberger, Matthew W. Self, Pieter R. Roelfsema, J. Alexander Heimel

## Abstract

Object detection is an essential function of the visual system. Although the visual cortex plays an important role in object detection, the superior colliculus can support detection when the visual cortex is ablated or silenced. Moreover, it has been shown that superficial layers of mouse SC (sSC) encode visual features of complex objects, and that this code is not inherited from the primary visual cortex. This suggests that mouse sSC may provide a significant contribution to complex object vision. Here, we use optogenetics to show that mouse sSC is causally involved in figure detection based on differences in figure contrast, orientation and phase. Additionally, our neural recordings show that in mouse sSC, image elements that belong to a figure elicit stronger activity than those same elements when they are part of the background. The discriminability of this neural code is higher for correct trials than incorrect trials. Our results provide new insight into the behavioral relevance of the visual processing that takes place in sSC.

## Introduction

When using vision to survey the environment, the brain segregates objects from each other and from the background. This object detection is an essential function of the visual system, since behavior typically needs to be performed in relation to the objects surrounding the organism. The primary visual cortex (V1) is involved in object segregation and detection (e.g. Lamme, 1995; Ress et al., 2000; Li et al., 2006). Silencing or ablating V1, however, does not completely abolish detection of simple visual stimuli in mice (Prusky & Douglas, 2004; Glickfeld et al., 2013; Resulaj et al., 2018; Kirchberger et al., 2021). (Glickfeld et al., 2013; Kirchberger et al., 2021; Prusky & Douglas, 2004)Furthermore, humans with bilateral lesions of V1 remain capable of detecting some visual stimuli, even in the absence of conscious vision; a phenomenon termed ‘blindsight’ (Ajina & Bridge, 2017). This capacity is likely to be mediated by the superior colliculus (SC), a sensorimotor hub in the midbrain (Tamietto et al., 2010; Kato et al., 2011; Ito & Feldheim, 2018; Kinoshita et al., 2019;

The SC receives direct input from the retina, as well as from V1 and other sensory areas (Basso & May, 2017), and mediates orienting responses to salient stimuli in primates and rodents (White & Munoz, 2012; Allen et al., 2021). In mice, detection of change in isolated visual stimuli is impaired in a spatial and time specific manner when the SC is locally and transiently inhibited (Wang et al., 2020). The mouse SC is also involved in hunting (Hoy et al., 2019; Shang et al., 2019) as well as defensive responses (Evans et al., 2018; Shang et al., 2018) to visual stimuli that are clearly isolated from the background. In recent years, a growing number of experiments point towards a role for the SC in more complex processes that are usually associated with the cerebral cortex (Krauzlis et al., 2013; Basso & May, 2017; Basso et al., 2021; Jun et al., 2021; Zhang et al., 2021). It is, however, not yet clear whether SC is also involved in detection of stimuli on a complex background.

When V1 is transiently silenced, mice are still able to detect stimuli that are defined by contrast on a homogeneous background, but they cannot detect texture-defined figures that only differ from the surrounding textured background by orientation (Kirchberger et al., 2021). Models for detection of texture-defined figure stimuli focus on the visual cortex and presume that interactions within V1 enhance the neural response to the figure edges, and that higher cortical visual areas feed back to ‘fill in’ the neuronal representation of the figure in V1 (Roelfsema et al., 2002; Poort et al., 2012; Liang et al., 2017). Indeed, neurons in V1 of mice and primates respond more vigorously to image elements in their receptive fields that differ from the surrounding texture (Lamme, 1995; Poort et al., 2012; Self et al., 2014; Li et al., 2018; Schnabel et al., 2018; Kirchberger et al., 2021). This figure-ground modulation (FGM) depends on feedback from higher visual cortical areas, just as was predicted by the models (Roelfsema et al., 2002; Keller et al., 2020; Pak et al., 2020; Kirchberger et al., 2021).

However, the ability to encode visual contextual effects is not exclusive to the visual cortex. The superficial layers of the rodent SC (sSC) have been shown to display orientation-tuned surround suppression (Girman & Lund, 2007; Ahmadlou et al., 2017; De Franceschi & Solomon, 2020): the responses of orientation-tuned neurons to an optimally oriented grating stimulus are attenuated when the surround contains a grating of the same orientation, whereas the responses are less suppressed or facilitated when the surround contains an orthogonal grating (Allman et al., 1985). This property is thought to play an important role in object segregation (Lamme, 1995). Interestingly, Ahmadlou et al. (2017) showed that the orientation-tuned surround suppression in sSC is computed independently of V1. The presence of this contextual modulation in the SC, combined with research showing the involvement of SC in visual detection (Wang et al., 2020) lead us to hypothesize that SC in the mouse might also be involved in detecting and segregating stimuli from a complex background. Here, we show that inhibiting the sSC reduces performance on a variety of figure detection tasks – indicating a causal role for the sSC in this behavior. Furthermore, we use extracellular recordings in mice performing figure detection to show that mouse SC indeed contains a neural code for figure detection.

## Results

### Superior colliculus is causally involved in figure detection

To test the involvement of superior colliculus in figure detection based on different features, we trained mice on three different versions of a figure detection task: a task based on figure contrast, a task based on figure orientation and a task based on figure phase (**Fig. 1A**). On each trial, the mice had to indicate the position of the figure (left vs. right) by licking the corresponding side of a Y-shaped lick spout (**Fig. 1B**). To test the involvement of the sSC in object detection, we optogenetically inhibited sSC while the mice performed the different tasks. The inhibition was achieved by injecting a viral vector with Cre-dependent ChR2, an excitatory opsin, in sSC of GAD2-Cre mice. We subsequently implanted optic fibers to target blue light onto the SC (**Fig. 1C**). Laser light activated the inhibitory neurons in sSC, thereby reducing the overall activity in superior colliculus (Ahmadlou et al., 2018; Hu et al., 2019). Anesthetized control experiments showed that the visually evoked responses in sSC were significantly reduced by 33% on average (p < 0.001), but we did not achieve complete silencing (**Fig. S1**). In order to test not only *if,* but also *when* superior colliculus is causally involved in figure detection, we inhibited the sSC at different latencies (0-200 ms) after stimulus onset. The mice were allowed to respond from 200 ms after stimulus onset (**Fig. 1D**). To prevent the behavioral report of the mice from being influenced by seeing the laser light turning on and off, we shielded the outside of the fiber implant. We also placed a blue LED above the head of the mouse that flashed at random intervals, which the animal learned to ignore. A total of N = 8 mice were used for these experiments. The mice typically performed 100-250 trials per session (one session per day, 5 sessions per week), and the recording period lasted for 2-5 months.

**Figure 1.**
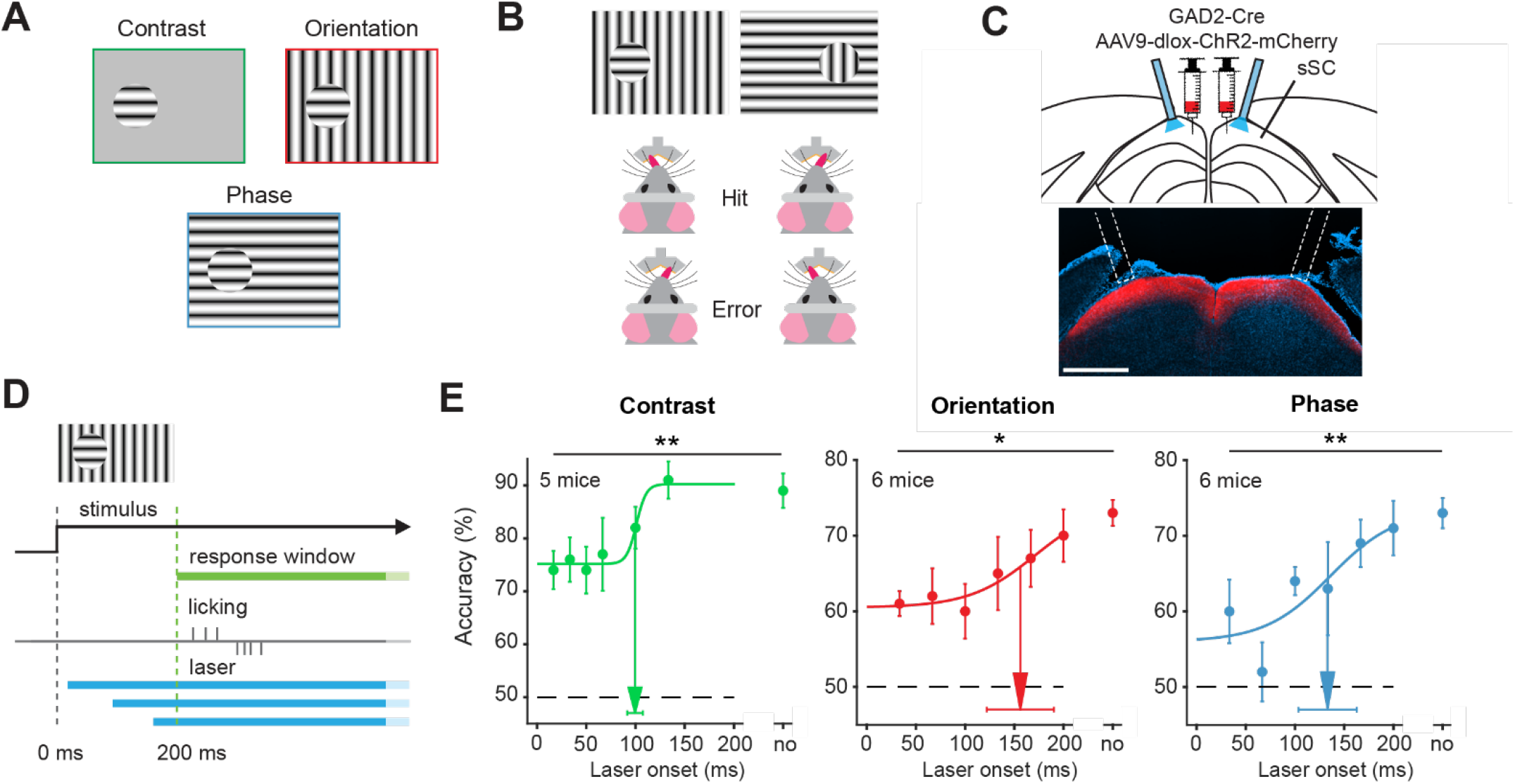
Superior colliculus is causally involved in figure detection. **(A)** The stimulus types that were used for the figure detection task. The stimulus consisted of a grating that differed from the background in either contrast (top left), orientation (top right) or phase (bottom). **(B)** Two example stimuli (both orientation task). Licking on the side corresponding to the figure constituted a hit, and vice versa. **(C)** Top: Schematic illustration of viral injections and optic fiber implantation. Bottom: histological verification of viral expression. Red: ChR2-mCherry. Blue: DAPI. **(D)** Timing of the task. We optogenetically inhibited activity in sSC in both hemispheres at different delays after stimulus appearance. The mice reported the figure location after 200 ms by licking on the same side as the stimulus. **(E)** Inhibition of sSC significantly decreased task performance for each figure detection task. The accuracy on unperturbed trials without the laser condition is indicated by ‘no’. Dots represent means ± SEM of accuracies across mice. Arrow and error bar indicate mean ± SD of bootstrapped fitted inflection points. Dashed line indicates chance level performance. *: p < 0.05, **: p < 0.01.

For each of the three figure detection tasks, inhibition of sSC significantly reduced the accuracy of the mice (**Fig. 1E,** p = 0.005, p = 0.027 and p = 0.003 for the contrast, orientation and phase task, respectively**. For details on all statistics in this paper, see Supp. Table 1**). This indicates that sSC is causally involved in figure detection, be it based on grating contrast, orientation or phase. In some mice, the optogenetic activation of inhibitory interneurons did not only decrease accuracy, but it also increased in the number of trials in which the mice did not respond at all (misses). When we also counted missed trials as error trials, the effect of the laser on the accuracy of mice was similar to the effects reported in **Figure 1** (**Figure S2,** p < 0.001, p = 0.24 and p = 0.002 for the contrast, orientation and phase task, respectively). Although the accuracies of the mice were decreased by the sSC inhibition, they typically were still above chance level. This might be related to the incomplete silencing of the sSC, although it is also possible that the thalamocortical pathway is involved in figure detection in parallel to the sSC. The accuracy of the mice recovered when we postponed the optogenetic inhibition. For the contrast task, the accuracy reached the half-maximal value when we postponed the laser onset to 99 ms (± 8 ms). In the orientation and phase task, the mice reached their half-maximum performance when we postponed the laser onset to 156 ms (± 35 ms) and 134 ms (± 30 ms), respectively. We therefore conclude that short-lasting activity in the sSC suffices for the contrast detection task, whereas orientation-and phase-defined object detection necessitate a longer phase of sSC activity. Based on comparisons between the response accuracy and processing time of the mice in their experiments, Kirchberger et al. (2021) concluded that it is not likely that these differences depend on task difficulty for the mouse. Rather, they probably reflect differences in the encoding strategy of the brain for the different tasks.

### sSC shows figure-ground modulation for contrast- and orientation-defined figures

Having confirmed that the sSC is causally involved when performing figure detection tasks, we next set out to investigate if neurons in the superior colliculus encode the visual information needed for the task, or encode decision-related information. Recent experiments have demonstrated that orientation- and phase-defined figures elicit more activity in the visual cortex of the mouse than the background image does; a phenomenon called figure-ground modulation (FGM) (Lamme, 1995; Kirchberger et al., 2021). To test whether this also occurs in sSC, we recorded neural activity in SC using 32-channel laminar electrodes while the mice performed the figure detection tasks (**Fig. 2A-B**). In this experiment, we kept the lick spout away from the mouse until 500 ms after stimulus onset to prevent noise caused by premature licks from interfering with the spike detection. After this delay, the lick spout automatically moved within licking distance (**Fig. 2C**). The mice could perform the tasks reliably above chance level during these recording sessions (**Fig. 2D**). During each recording session, we first mapped the receptive fields (RFs) of the recorded sites. We then set up the visual stimuli of the figure detection task such that in each trial, the figure stimulus was placed either over the RF, or 50-60 degrees lateral from the RF in the other visual hemifield (**Fig. 2E-F**). This way, as the mouse was reporting the side of the figure stimulus in each trial, the recorded neurons would variably respond to the figure or to the background. By varying the orientations of the stimuli, we could compare the figure and background condition using trials that had identical information presented inside the receptive field. We recorded neural activity from 5 mice.

**Figure 2.**
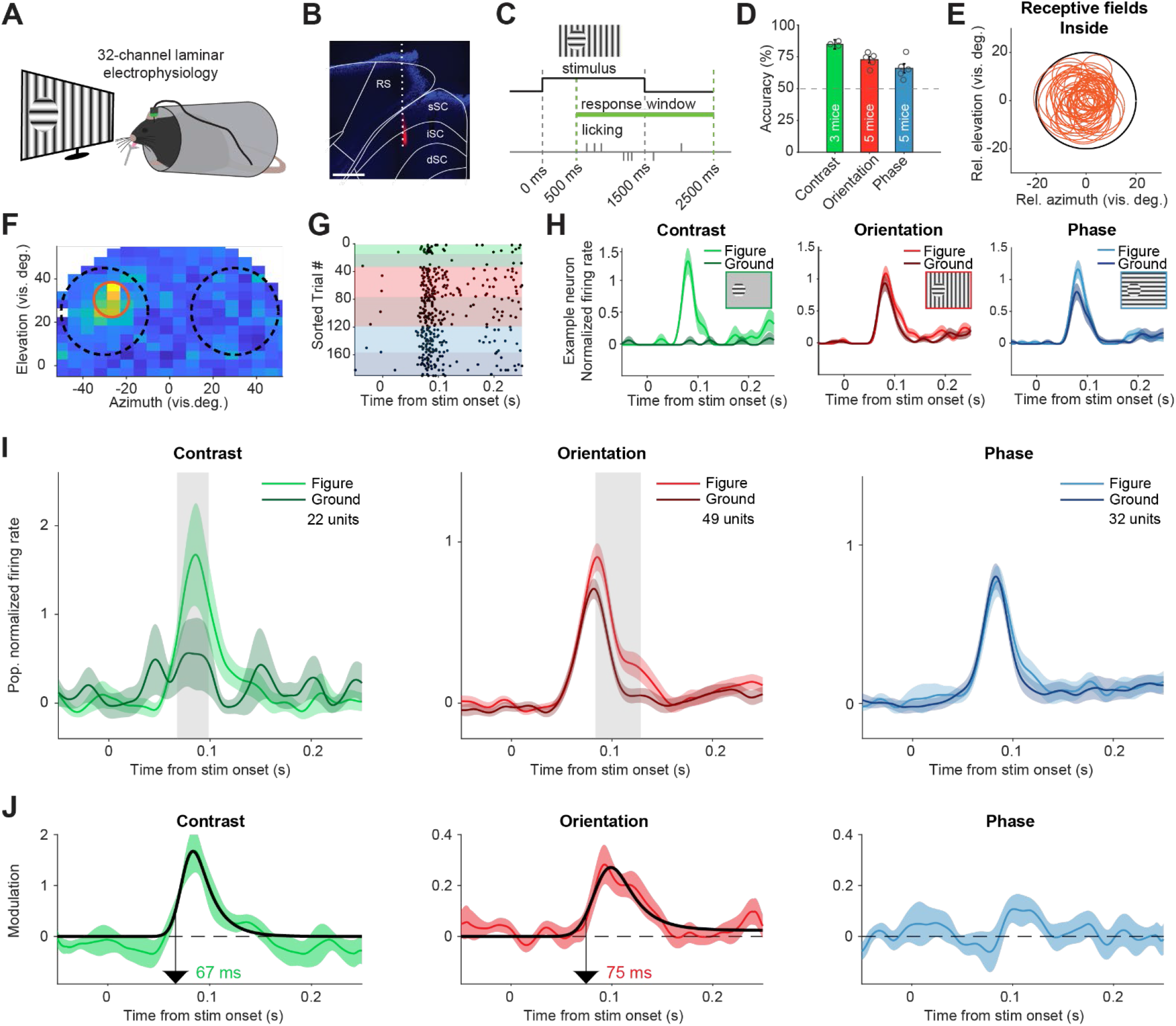
Superior colliculus activity elicited by contrast and figure-ground stimuli. **(A)** Schematic illustration of setup. **(B)** Histological verification of electrode track. Blue: DAPI. Red: diI. **(C)** Timing of the task. The mice could report the figure location after 500 ms. **(D)** Accuracy of the mice in each task, mean ± SEM. **(E)** Estimated receptive fields of neurons with an RF entirely inside the figure. **(F-H)** Example neuron. **(F)** Receptive field of an example neuron. Black circles indicate the position of the figure (on top of RF) and ground (outside of RF) stimulus in the visual field. Red circle indicates estimated RF. **(G)** Raster plot, sorted by task and trial type. Each dot indicates a spike. Green, red and blue colors indicate contrast, orientation and phase task trials, respectively. Brighter colors indicate figure trials, darker colors indicate ground trials. **(H)** Mean (± SEM) activity of the example neuron for each task. **(I)** Mean (± SEM) population responses for each task. Grey patches indicate time clusters where the difference between figure and ground is significant (p < 0.05). **(J)** Difference between figure and ground responses in each task and estimated onset of the response difference. Colored lines indicate data, black lines indicate fit of the response. Arrows indicate the onset latency of the response difference.

In our analysis, we only included neurons with an estimated RF completely inside of the figure stimulus (**Fig. 2E**). The contrast-defined figure elicited a strong response compared to the gray background, with an estimated onset of 67 ms (**Fig. 2H-J**, left; p < 0.05 from 68-99 ms in Fig. 2I). For the orientation-define figures we also found significant FGM, with an estimated onset of 75 ms (**Fig. 2H-J**, middle; p < 0.05 from 84-129 ms in Fig. 2I). When the figure was defined by a phase difference, FGM was not significant (**Fig. 2H-J**, right; p > 0.05 for all time bins in Fig. 2I). We conclude that the activity of sSC neurons elicited by contrast- and orientation-based figures is stronger than that elicited by a background. This effect did not occur for phase-defined figures, even though optogenetic sSC inhibition had impaired the accuracy of the mice in the task in which they had to locate phase-defined figures.

### Superior colliculus represents the location of figures

Our data indicates that phase-defined figures on average elicit a similar response as the background. However, the task performance of the mice did decrease when inhibiting the sSC during the phase task. We therefore examined the neuronal responses in more detail. Whereas the results in **Figure 2** represented neurons with RFs confined to the figure interior, we also recorded neurons with RFs on the edge of the figure stimulus (**Fig 3A**). For these neurons, we found a significantly higher response for figure vs. ground in the orientation task, but not in the phase task (**Fig. 3B**; p < 0.05 from 81-102 ms in the orientation task, p > 0.05 for all time bins in the phase task). We did not record enough edge-RF data from the contrast task so it is excluded here.

**Figure 3.**
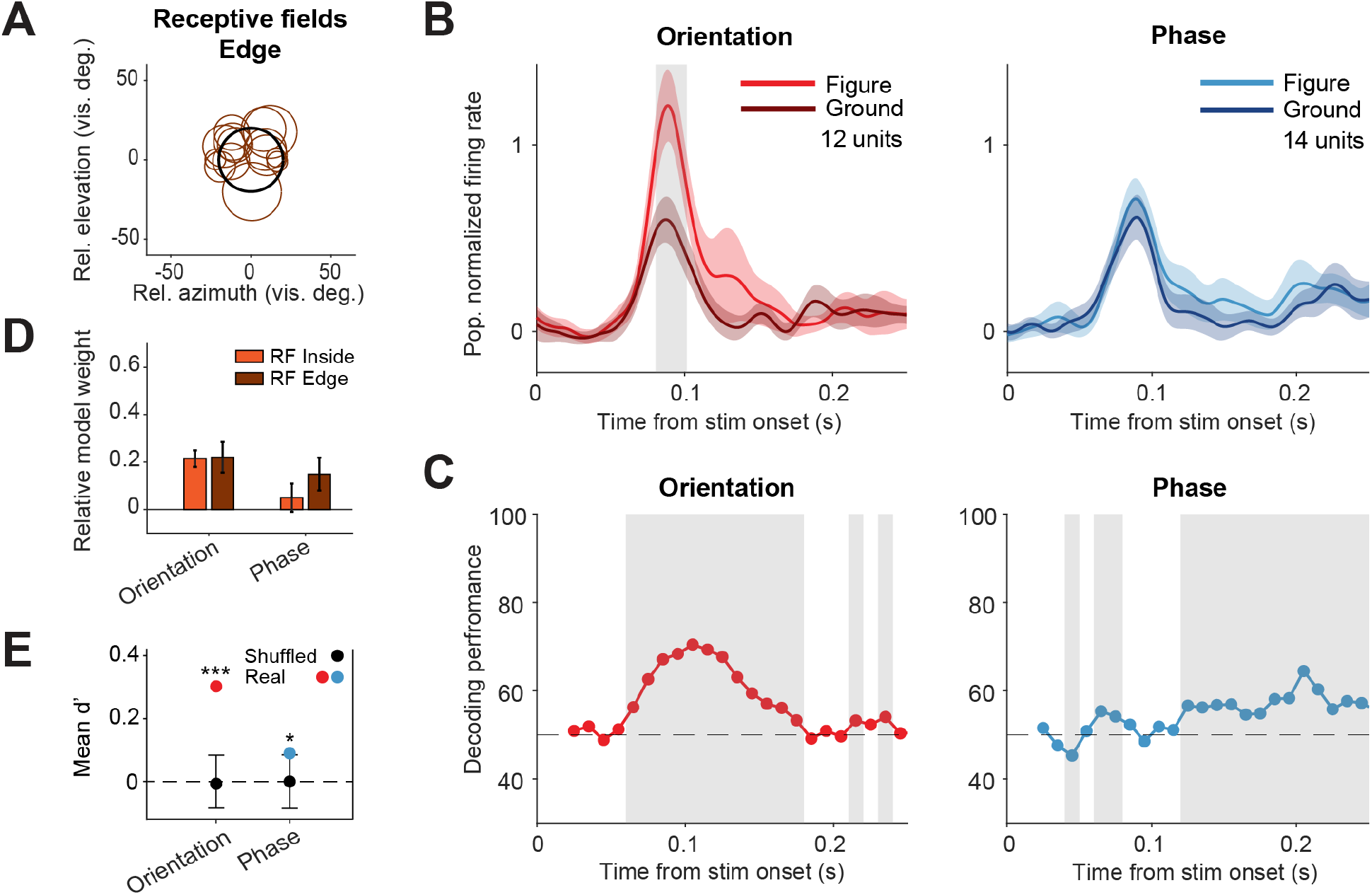
Decoding the stimulus identity from population responses in superior colliculus. **(A)** Estimated receptive fields of the neurons with an RF on the figure edge. **(B)** Mean (± SEM) population responses of the RF edge-neurons for each task. Grey patches indicate time clusters where the difference between figure and ground is significant (p < 0.05). **(C)** Decoding performance of a linear SVM classifier for each task. Performance was computed using a sliding window of 50 ms in steps of 10 ms. Grey regions indicate decoding performance significantly different from chance (p<0.05). **(D)** Mean (± SEM) relative model weights for each task, comparing weights of RF-inside neurons with those of RF-edge neurons **(E)** Neuronal d-primes during the window with the best decoding performance. Black bars indicate the mean (± 95% confidence interval) population d-primes of bootstraps with shuffled trial identities. Colored lines indicate real mean of d-primes in the population. *: p<0.05, **: p<0.01, ***: p<0.001.

We conclude that the phase-defined figure-ground stimulus did not cause a significantly enhanced population response in the sSC. It is, however, conceivable that the figure could be represented by some neurons that enhance their response and others that decrease their response, without an overall influence of the firing rate at the population level. To examine this possibility, we used a linear SVM to decode the stimulus identity (figure vs. ground) from the recorded population, including both the RF-inside and RF-edge neurons. As the data set stems from mouse behavior, it was not balanced with regard to trial numbers per trial type. In order to correct for this, we used a bootstrapping strategy (see methods section). In brief, we trained the model many times, each time on a different balanced pseudo-randomized sub-selection of the trials, and using leave-one-out cross-validation to test the model performance (**Fig. 3C**). For the orientation task, the decoder could detect the stimulus identity at a performance of around 75% (lowest p < 0.001 at time window 80-130 ms). Interestingly, the decoder also detected the phase stimulus identity above chance level, more specifically in later time windows, after the peak of the visual response (lowest p < 0.001 at time window 180-230 ms). These results are in line with the onset and relative strength of the figure-ground response difference for each task, and show that visual information for performing the task is present in the sSC.

To understand which information was used by the SVM model, we first analyzed the model weights of the individual neurons, split by RF type (**Fig. 3C**). For the orientation task, the relative weights of the RF-center and RF-edge neurons were very similar (p = 0.863), indicating that the responses of those groups of neurons contain roughly equal information about the stimulus. For the phase task, the RF-edge neurons had higher relative weights than the RF-center neurons, although the difference was non-significant (p = 0.277). Higher relative weights for the RF-edge neurons would suggest that sSC encoded the phase-defined figure mainly by using edge-detection.

We also computed the d-prime of the recorded neurons; a measure of the reliability of the difference between the figure and ground response on single trials (see methods section). For both tasks, the d-primes were significantly higher than chance level in the time window with the best SVM decoding performance (**Fig. 3E**; p < 0.001 and p < 0.05 for orientation and phase, respectively). We conclude that although the difference between figure vs. ground responses was small in the phase task, the variability of the neuronal responses was low enough for reliable decoding.

### Different discriminability in sSC preceding hits vs. errors

Because superior colliculus is a sensorimotor hub, we wanted to further investigate whether the colliculus might not only encode the visual stimulus but also the decision of the mouse. To this end, we split up the data from the visually responsive neurons between hit (correct) and error (incorrect) trials (**Fig. 4A**), and analyzed the firing rates between stimulus onset and the response of the mouse. We did not have enough data to perform this analysis for the contrast task, and we therefore focus on the orientation and phase tasks. The figure-ground modulation in sSC was more pronounced on hit trials than on error trials, both for the orientation task and the phase task (**Fig. 4B**). To statistically compare the response difference between hit and error trials, we computed d-primes and created a mixed linear effects model (**Fig. 4C**). The discriminability (d-prime) was significantly higher for hit trials than error trials (p < 0.001), showing that activity in the sSC is reflected in the behavioral performance. Post-hoc analysis showed that the difference in discriminability was mainly driven by the difference between hits and errors in the orientation task (**Supp Table 1;** p < 0.001 and p = 0.303 for orientation and phase, respectively).

**Figure 4.**
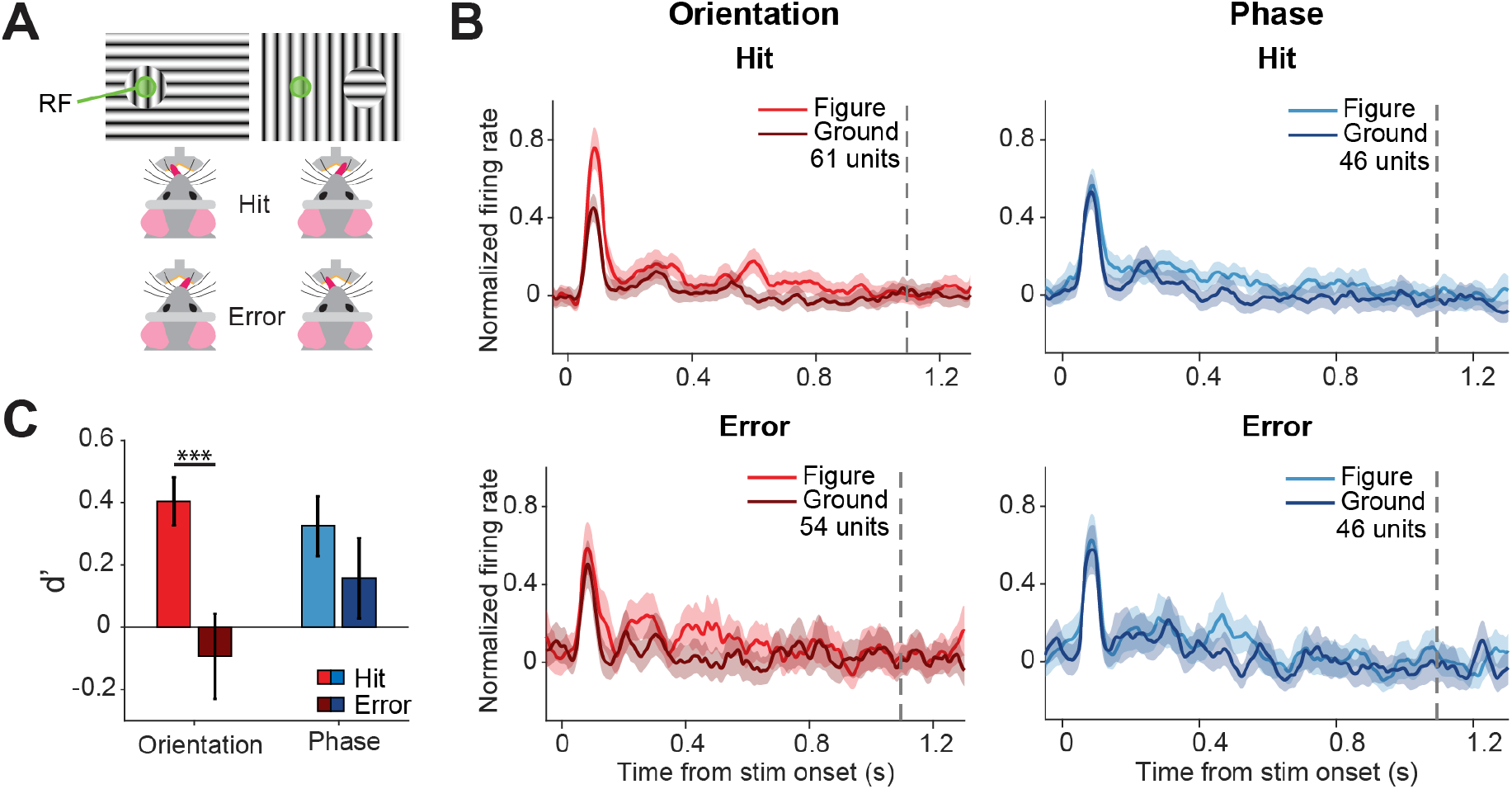
Different discriminability in sSC for hits vs. errors. **(A)** Two example stimuli (both orientation task). Licking on the side corresponding to the figure constituted a hit, and vice versa. RF: receptive field. Note that the information inside the receptive field is the same between the two stimuli. **(B)** Population responses for hits vs. errors. Dashed grey line indicates the mean reaction time of the mice. Note that the difference between figure and ground is larger for hits than errors. **(C)** Neuronal d-primes were higher for hit trials than for error trials. ***: p<0.001

### A group of putative multisensory neurons encodes both the stimulus and the decision

Not all neurons in our data set were exclusively visually responsive. A small group of neurons only reached their peak firing rates around 700 ms after stimulus onset (**Fig. 5A**). Although late, these responses were time-locked to the onset of the trial (p<0.05 Zeta-test for all cells; Montijn et al., 2021). These neurons presumably responded to the movement of the lick spout towards the mouse after 500 ms, providing auditory and somatosensory input (the spout moved below the head of the mouse and therefore wasn’t visible to the mouse). The activity of these neurons did not correlate with eye movements (**Fig. S3A-B;** p = 0.488). Furthermore, their sharply timed firing rate peak is not present when plotting the responses relative to the time of the lick, indicating that these neurons are probably not directly involved in movement initiation (**Fig S3C**). Given their putative response to the spout movement, combined with their modest response to the visual stimulus, we call these neurons putative multisensory neurons.

**Figure 5.**
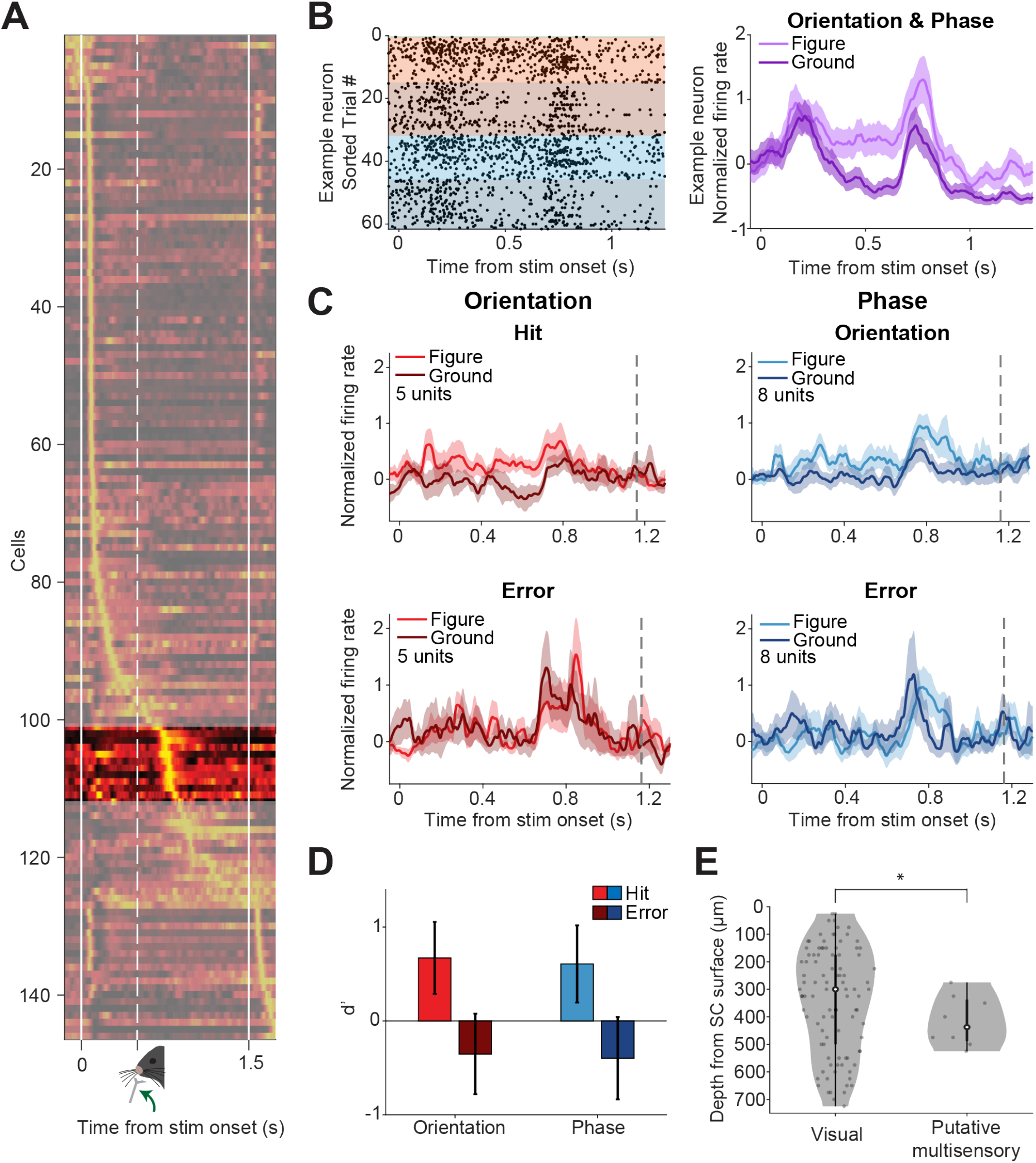
A group of putative multisensory neurons shows task-related responses. **(A)** Mean responses of all recorded neurons, sorted by the time of their peak response. A small group of neurons (region without shading) is most active around 700 ms: just after the lick spout moves towards the mouse, but before the mouse licks. **(B)** Responses of example putative multisensory neuron. Left: Raster plot where each dot indicates a spike. Red and blue indicate orientation and phase task trials, respectively. Brighter colors indicate figure trials, darker colors indicate ground trials. Right: Mean response of example neuron for figure vs. ground trials (orientation and phase trials combined). Shading indicates SEM. **(C)** Population figure-ground responses for hits and errors across all putative multisensory cells. Dashed grey line indicates the mean reaction time of the mice. Note that the response difference between figure and ground trials is larger for hits than for errors. **(D)** Neuronal d-primes for hit and error trials. **(E)** Comparison of estimated histological depth of visual vs. putative multisensory cells. The variances of the two groups are different. *: p<0.05.

As multisensory neurons in SC are generally found in deeper layers than visual neurons, we compared the estimated histological depths of the putative multisensory neurons with those of the visually responsive neurons (**Fig. 5E**). The putative multisensory neurons had a smaller variation in histological depth than the visual neurons (p = 0.046), suggesting that they represent a different cell population. When examining the responses of the putative multisensory neurons we found that, despite not being primarily responsive to the onset of the visual stimulus, they did respond more strongly to figure stimuli than ground stimuli (**Fig. 5B-C**). Just like the visual neurons, the d-primes of the putative multisensory neurons were significantly lower for error trials than hit trials (**Fig. 5D**; p = 0.044 for the main effect of response, post-hoc analysis showed p > 0.05 for the individual tasks).

## Discussion

Our experiments show that the superficial layers of superior colliculus are causally involved in detecting objects on a non-homogeneous background and detecting objects based on figure contrast, orientation and phase. Indeed, neurons in sSC show an increased response to figure stimuli compared to ground stimuli in both the contrast and orientation task. We did not find a significantly increased population response for phase-defined figures, but a linear SVM decoder indicated that the response difference was consistent enough to decode the phase stimulus above chance level. The discriminability between figure- and ground responses was higher for hit trials than error trials, suggesting that sSC may contribute to the decision of the mouse. Finally, we recorded a small group of putative multisensory neurons. These showed task-related responses similar to the visual neurons, but their activity peaked at a later point in time during the trial.

This study was aimed at the superficial part of SC (i.e. the stratum zonale, stratum griseum superficiale and the stratum opticum), although we cannot rule out the possibility that in some animals deeper parts of SC were recorded or were affected by optogenetics. Interestingly, to our knowledge this is one of only few studies that specifically inhibited the superficial part of SC bilaterally. Many previous studies have inhibited or ablated superior colliculus, but often this was done in a unilateral fashion and/or targeting the deeper layers or the entirety of the colliculi (Tunkl & Berkley, 1977; Mohler & Wurtz, 1977; Tan et al., 2011; Wolf et al., 2015; Ahmadlou et al., 2018; Hu et al., 2019; Wang et al., 2020; but see also Casagrande & Diamond, 1974). Therefore, our study provides new insight in the behavioral relevance of the visual processing that takes place specifically in sSC. The few putative multisensory neurons that we recorded at a slightly deeper location give initial insight in how task-related information is processed in downstream parts of the colliculus. Optogenetic inhibition was done by activating GABA-ergic neurons in the sSC. This is a commonly used strategy to inhibit areas in the neocortex (Lien & Scanziani, 2018; Vangeneugden et al., 2019), where GABA-ergic neurons only project locally. It allows relatively long and repeated silencing without unwanted side-effects. Most of the GABA-ergic neurons in the sSC are also locally projecting, but there is a small fraction of GABA-ergic neurons projecting out of the superior colliculus (Gale & Murphy, 2014; Whyland et al., 2020; C. Li et al., 2022) that hypothetically could directly mediate the effects of the sSC on behavior.

Our results suggest that superior colliculus is necessary not just for simple but also for relatively complex object detection, when the figure does not stand out from the background by contrast. Previous experiments have already shown that the SC is causally involved in detecting orientation change (Wang et al., 2020), looming stimuli (Evans et al., 2018; Shang et al., 2018) and detecting moving objects during hunting (Hoy et al., 2019). Here, we show that SC is also involved in the detection of more complex, static objects. This behavior is often called figure-ground segregation, but we have to point out an important difference between previous figure-ground segregation research (e.g. Lamme, 1995; Poort et al., 2012; Jones et al., 2015), and our study. First, unlike commonly used stimuli for macaques, our stimuli showed a clear figure edge, due to the adaptation of the stimulus to mouse acuity. In addition to that, the tasks did not involve any eye fixation. Therefore, it would have been a viable task strategy for the mouse to simply inspect the figure edge – a strategy that is normally prevented by the high and varied spatial frequency of the stimuli for macaques. So although we can be sure that the mice perform object *detection,* we cannot be entirely sure that they perform object *segregation* in the purest sense of the word. Because our task used a limited number of grating orientations and positions, a potential task strategy would be for the mice to learn the correct responses to the complete image of figure and background. Earlier experiments with the same stimuli in freely walking mice, however, suggested that after training on a restricted training set, mice generalize over size, location and orientation of the figure gratings, and therefore perform the task as if they detect the presence of an object (Schnabel et al., 2018). In any case, our results add to the growing number of experiments (Basso et al., 2021; Bogadhi & Hafed, 2022) that nuance and expand upon the view of the superior colliculus as a saliency map that depends on the visual cortex for complex visual processing (Lamme et al., 1998; Fecteau & Munoz, 2006; Gilbert & Li, 2013; Zhaoping, 2016; White et al., 2017).

Interestingly, the accuracy of the mice did not decrease to chance level when inhibiting the sSC. This might be partially due to the incomplete silencing of sSC (**Fig. S1**), but may also suggest that other brain areas contribute in parallel to the performance of this visual task. The obvious candidate area for this is the visual cortex. Many studies have shown the involvement of mouse V1 in object detection (Glickfeld et al., 2013; Katzner et al., 2019). Evidence that SC works in parallel to the visual cortex comes from the finding that mice can perform the contrast detection task above chance level when V1 is silenced (Kirchberger et al., 2021). The detection of orientation-defined and phase-defined figures, however, was abolished when V1 was silenced (Kirchberger et al., 2021). This suggests that the parallel processing stream through the superior colliculus does not suffice for detection of these more complex figures.

To increase our understanding of the role division between sSC and V1 in object detection, we can compare our sSC results to the V1 results from Kirchberger et al. (2021). When inhibiting sSC during object detection, we estimated the half-maximum performance times to be 99 ms, 156 ms, and 134 ms for detection based on contrast, orientation, and phase, respectively. In V1, the corresponding half-maximum performance times were 62 ms, 101 ms and 141 ms. The parsimonious interpretation of these findings is that V1 performs a role in object detection at an earlier point in time than sSC (for object detection based on contrast and orientation, but not phase). From there, we could hypothesize that sSC inherits some of its task-related code from V1 and that sSC operates downstream of V1 in the object detection process. Indeed, superior colliculus is involved in sensorimotor transformations (e.g. Gandhi & Katnani, 2011; Duan et al., 2021). However, the onset times of the neural responses paint a slightly different picture: in V1, the onset times of the FGM for contrast, orientation and phase were estimated at 43 ms, 75 ms and 91 ms, respectively. The onset times we recorded in sSC were 67 and 75 ms for contrast and orientation stimuli, and a non-significant result for the phase stimuli. Although our estimates of the onset time of FGM are not precise enough to fully rule out the possibility that modulation of V1 is transferred to sSC through the direct projection, the onset time of the modulation in sSC suggests that it is computed independently of V1. This independency of V1 has also been shown for orientation-dependent surround suppression in sSC (Girman & Lund, 2007), which even increases if V1 is inhibited (Ahmadlou et al., 2017). This suggests a role for the sSC upstream of V1 or in parallel to V1, perhaps in strengthening the representation of pop-out stimuli in V1 through pathways that include LP (Hu et al., 2019; Fang et al., 2020) or LGN (Jones et al., 2015; Ahmadlou et al., 2018; Poltoratski et al., 2019).

Our study provides evidence for a neural code in mouse sSC that is necessary for normal visual detection of complex static objects. These results fit in with the growing number of studies that show mouse sSC provides a significant contribution to the processing of complex visual stimuli (Ahmadlou et al., 2017; Hu et al., 2019; Fang et al., 2020). The superior colliculus (or the optic tectum in non-mammalian species) is a brain area conserved across vertebrate evolution (Isa et al., 2021). Even though clear differences exist between mouse and primate sSC, for example in their received retinal inputs (Ito & Feldheim, 2018), representation of visual features (Wang et al., 2010; Ahmadlou & Heimel, 2015; Chen & Hafed, 2018), and more generally the animals’ strategies for visual segmentation (Luongo et al., 2021), many functions of sSC are shared between rodents and primates. Some of these include saliency mapping (White et al., 2017; Barchini et al., 2018), spatial attention (Krauzlis et al., 2013; Wang & Krauzlis, 2018; Hu & Dan, 2022; Wang et al., 2022) and orienting (Boehnke & Munoz, 2008; Masullo et al., 2019; Zahler et al., 2021). This and recent work showing object coding in the primate (Griggs et al., 2018; Bogadhi & Hafed, 2022) suggests that also primate sSC contributes to visual processing in more various and complex ways than anticipated based on previous work.

## Materials & Methods

All offline analysis was performed using MATLAB (R2019a; MathWorks).

### Experimental animals

For the experiments we used a total of 16 mice. For awake behaving electrophysiology, we used 5 C57BL/6J mice (Charles River, all male). For behavior combined with optogenetics, we used 8 GAD2-Cre mice (Stock #028867, Jackson; 6 male, 2 female). For anaesthetized electrophysiology, we used 3 GAD2-Cre mice (all male). Mice were 2-5 months old at the start of experiments. The mice were housed in a reversed light/dark cycle (12 hr / 12 hr) with ad libitum access to laboratory food pellets. All experiments took place during the dark cycle of the animals. Mice were either housed solitarily or in pairs. All experimental protocols were approved by the institutional animal care and use committee of the Royal Netherlands Academy of Sciences (KNAW) and were in accordance with the Dutch Law on Animal Experimentation.

### General surgical preparation and anesthesia

Anesthesia was induced using 3-5% isoflurane in an induction box and was maintained using 1.2-2% isoflurane in an oxygen-enriched air mixture (50% air and 50% O_2_, 0.5 L per min flow rate). After induction, the mice were positioned in a Kopf stereotactic frame. The temperature of the animal was monitored and kept between 36.5° and 37.5° using a heating pad coupled to a rectal thermometer. We subcutaneously injected 2.5 mg/kg meloxicam as general analgesic, and the eyes were covered with Bepanthen ointment to prevent dehydration and to prevent microscope light from entering the eye. The depth of anesthesia was monitored by frequently checking paw reflexes and breathing rate. We added a thin layer of xylocaine cream to the ear bars for analgesia and stabilized the mouse’s head in the frame using the ear bars. Then the area of the incision was trimmed or shaved, cleaned with betadine, and lidocaine spray was applied to the skin as a local analgesic. We made an incision in the skin, and then again applied lidocaine, this time on the periosteum. Further methods for specific surgeries are described below. At the end of each surgery, we injected 2.5 mg/kg meloxicam for post-surgical analgesia and kept the mice warm until they had woken up. We monitored their appearance and weight daily for at least 2 days post-surgery.

### Head bar implantation

After induction of anesthesia as described above, we cleaned the skull, thereby removing the periosteum, and slightly etched the skull using a micro curette. We then applied a light-cured dental primer (Kerr Optibond) to improve the bonding of cement to the skull. After applying the primer, we created a base layer of cement on top of the primer using Heraeus Charisma light-cured dental cement. The head bar was placed on top of this base layer and fixed in place using Vivadent Tetric evoflow light-cured dental cement. Lastly, we sutured the skin around the implant.

### Viral injections

We diluted ssAAV-9/2-hEF1α-dlox-hChR2(H134R)_mCherry(rev)-dlox-WPRE-hGHp(A) (titer 5.4 x 10^12^ vg/ml, VVF ETH Zurich) 1:1 in sterile saline and loaded it into a Nanoject II or Nanoject III injector (Drummond Scientific). After induction of anesthesia as described above, we drilled two small craniotomies (0.5 mm in diameter) bilaterally above superior colliculus (0.3 mm anterior and 0.5 mm lateral to the lambda cranial landmark). Next, we inserted the pipette and slowly injected the viral vector solution at 2 different depths (55 nl each at 1.4 and 1.2 mm depth). After each depth, we waited 2 minutes before moving the pipette up. We left the pipette in place for at least 10 min before fully retracting it to avoid efflux. We repeated this for the second hemisphere. After the injections, we cleaned the scalp with sterile saline and sutured the skin.

### Fiber implant surgery

Optic fiber implants were custom made using grooved ferrules (Kientec Systems Inc., ID 230um, L=6.45mm, OD=1.249mm), multimode optic fiber (Thorlabs FP200URT, NA=0.5), and 2-component epoxy glue. Fiber implant surgery was performed at least one week after viral injection.

After induction of anesthesia as described above, we cleaned and etched the skull using sterile saline and a micro curette. We then applied a light-cured dental primer (Kerr Optibond) to improve the bonding of cement to the skull. Then, we drilled two small craniotomies (0.5 mm in diameter) above bilateral superior colliculus (0.5 mm anterior and 0.8 mm lateral from the lambda cranial landmark). We put an optic fiber in a custom holder at a 14° angle in the mediolateral plane – the angle prevented the fibers from blocking each other’s connection sleeve – and inserted it 0.9 mm deep. We added Vivadent Tetric evoflow light-cured dental cement to stabilize the implant and then removed the fiber from the holder. This was repeated for the second hemisphere. Once the optic fibers were thoroughly stabilized with cement, we placed a head bar anterior of the optic fibers, as described above. After this we sutured the skin around the implant.

### Surgery for awake electrophysiology

After mice had learned the task, we performed a surgery in which we made a craniotomy and placed a reference screw to enable awake behaving electrophysiology. After induction of anesthesia and before opening the skin, we injected 3 mg/kg dexamethasone s.c. to prevent cortical edema. We made an incision in the skin over the midline, posterior to the head bar implant. With a small razor, we cut the tendons of the neck muscle on the occipital bone to create space on the bone. After this we cleaned and dried the skull and applied light-cured dental primer (Kerr Optibond) for adhesion. We marked the location of the center of the craniotomy for the electrode insertion (0.5 mm anterior and 0.5 mm lateral from the lambda cranial landmark). We then first drilled a 0.6 mm craniotomy for the reference screw in the occipital or parietal bone, contralateral of the craniotomy, and inserted the screw. For some mice, we repeated this for a second screw to separate the electrical reference and ground. We then used Vivadent Tetric evoflow light-cured dental cement to stabilize the screws on the skull and to create a well around the marked location for the craniotomy. This well could hold a bit of sterile saline during the recordings to prevent desiccation of the brain tissue. We then continued to drill a 1.5-2 mm craniotomy above SC. We thoroughly cleaned the craniotomy with sterile saline and used sterile silicone (Kwik-Cast, World Precision Instruments) to seal the well. Finally, we sutured the skin around the implant.

### Visual stimulation

#### Visual stimulation in behavior task

For all the experiments including behavior, we created the visual stimuli with the Cogent toolbox (developed by J. Romaya at the LON (Laboratory of Neurobiology) at the Wellcome Department of Imaging Neuroscience) and linearized the luminance profile of the monitor/projector. For the optogenetic experiments, we used a 23-inch LCD monitor (1920 × 1080 pixels, Iiyama ProLite SB2380HS), placed 12 cm in front of the eyes. For the behavioral electrophysiological experiments, the stimuli were presented on a 21 inch LCD monitor (1280 × 720 pixels, Dell 059DJP) placed of 15 cm in front of the mouse. Both screens had a refresh rate of 60Hz. We applied a previously described correction (Marshel *et al.,* 2011) for the larger distance between the screen and the mouse at higher eccentricities. This method defines stimuli on a sphere and calculates the projection onto a flat surface. The figure and background were composed of 100% contrast sinusoidal gratings with a spatial frequency of 0.08-0.1 cycles/deg and a mean luminance of 20 cd/m^2^. The diameter of the figure was 35° (optogenetics) or 40° (electrophysiology). For the contrast-defined stimuli, we presented the figure gratings on a gray background (20 cd/m^2^). For the orientation-defined figures, the grating orientation in the background was either horizontal or vertical (0° or 90°), and the orientation of the figure was orthogonal. For the phase-defined figures, the phase of the figure grating was shifted by 180° relative to that of the background.

#### Visual stimulation for anesthetized experiments

During anesthetized electrophysiological experiments, visual stimuli were projected onto a back-projection screen placed 15 cm from the mouse with a PLUS U2-X1130 DLP projector (mean luminance = 40.6 cd/m^2^). The size of the projection was 76 cm by 56 cm, the field-of-view was 136° × 101.6°, the resolution was 1024 × 768 pixels, and the refresh rate was 60 Hz.

### Behavioral task

The mice were handled for 5 to 10 min per day for at least 5 days before training them in the setup. For the behavior sessions, the mice were head restricted in a tube. We habituated the mice to the head restriction by putting them in the setup each day for at least 5 days, ramping up the time in the setup from several minutes to ca. 30 minutes. Once they were fully habituated, they were put on a fluid restriction protocol with a minimal intake of 0.025 ml/g per day (in line with national guidelines, i.e. www.ncadierproevenbeleid.nl/adviezen-ncad), while their health was carefully monitored. The minimal intake was guaranteed by monitoring the water intake during the task performance. If the intake after behavioral experiments was below the daily minimum, mice were given HydroGel (Clear H_2_O) to reach the minimum.

The animals were trained to indicate the side on which a figure appeared by licking the corresponding side of a custom-made y-shaped lick spout (**Fig. 1B**). We registered licks by measuring a change in either capacitance (for optogenetics experiments) or current (for electrophysiology experiments) with an Arduino and custom-written software. A trial started when the stimulus with a figure on the left or right appeared on the screen. The stimulus was displayed for 1.5 s. Because mice made early random licks, we disregarded licks from 0 to 200 ms. The exact figure location varied slightly depending on the RF positions, but the figure center was generally close to an azimuth of 30° (left or right of the mouse) and an elevation of 15°. Correct responses were rewarded with a drop of water. Stimulus presentation was followed by a variable intertrial interval (ITI) of 6 to 10 s. If the animal made an error, a 5-s timeout was added to the ITI. We presented a background texture during the ITI and did not give reward if the mice licked so that they learned to ignore the background. In some sessions, we included correction trials, which were repeats of the same trial type after an error. We only included non-correction trials for our analysis of the accuracy of the mice. During the electrophysiology experiments, we used an Arduino-controlled servo motor that moved the lick spout towards the mouth of the mouse 500 ms after the presentation of the stimulus, thereby ensuring that the first 500 ms of the visual response could be recorded without electrical artefacts from the lick detector. We define task accuracy as hits/(hits + errors). For **Fig. S2** only, accuracy was defined as hits/(hits + errors + misses).

The training to the task involved various steps of increasing difficulty. First, the mice were trained on the contrast task, detecting the position of a circular grating figure on a grey (20 cd/m^2^) background. After learning this, the mice were introduced to different backgrounds. This was done by starting out with a figure grating on a black (0 cd/m^2^) or white background (40 cd/m^2^). Essentially this meant that the contrast of the grating in the background was 0%. When the mouse performed above 70% accuracy, the background grating would gradually change from 0% contrast to 100% contrast. Likewise, when the mouse performed below 60% accuracy, the background grating would decrease in contrast. When they reached 100% contrast, the mice had learned the orientation task. We then introduced phase-defined stimuli in 50% of the trials so they could generalize to these stimuli.

### Optogenetic inhibition during behavior

For bilateral optogenetic inhibition of SC through activation of GABA-ergic neurons, we used either a 473 nm BL473T8-100FC laser or a 462 nm BLM462TA-100F laser (Shanghai Laser & Optics Century Co.). The laser was connected to a two-way split patch cord (Doric Lenses), with a power of 3-5 mW at each fiber tip. We placed a blue distractor LED light, driven by an Arduino, at a height of 38 cm above the mouse. The light flickered on for 0 – 1.2 seconds and off for 0 – 2.8 seconds to habituate the mouse to any flickering stray blue light. The contrast task was recorded in sessions that were independent from the sessions with the orientation- and phase-defined task. In the orientation/phase sessions, orientation-defined and phase-defined stimuli were pseudorandomly shuffled in a 50/50% ratio. The mouse was placed in the setup, and the patch cord was connected to the bilateral implant. The implant was shielded using Creall super soft modelling clay to prevent light scatter out of the implant. When the animals performed the task consistently with an accuracy larger than 65%, we initiated the inhibition of SC using laser light in a random 25% of the trials. The onset of stimulation was shifted relative to the onset of the visual stimulus in steps of 16.7 ms, conforming to the frame rate of the screen. The optogenetic stimulation lasted for 2 seconds.

For the analysis, we included only trials from the periods in recording sessions where optogenetic inhibition was used, i.e. periods in which the mice had an accuracy higher than 65%. In our analysis of the influence of optogenetic silencing, we computed the accuracy for each laser onset latency for each mouse. We fit a logistic function to the mean accuracies using the Palamedes toolbox in MATLAB (Prins & Kingdom, 2018). In order to get a good estimate of the time at which the accuracy reached its half maximum (i.e. the inflection point of the fitted curve), we used bootstrapping (1000 times) by sampling trials from each mouse with replacement. For each bootstrap, we fit a logistic function to the results, resulting in a distribution of estimated inflection points. To test the significance of the effect of the optogenetic inhibition on the accuracy of the mice, we used one-way repeated measures ANOVA with Bonferroni correction.

### Awake behaving electrophysiology

After the craniotomy surgery and at least 2 days of recovery, the mice were recorded daily for up to two weeks. First the mouse was placed in the setup. The left eye of the mouse was tracked using an ISCAN camera and ISCAN software. We used matte black aluminum foil (Thorlabs) to shield its eyes from light during electrode insertion, and also to shield the craniotomy from electrical noise. During the last recording session of each mouse, we coated the electrode tip with diI for histological verification. While looking through a microscope (Zeiss Stemi 508), we removed the Kwik-Cast from the well and cleaned the recording chamber with sterile saline. We connected the ground/reference screws to the recording system, and slowly inserted a Neuronexus probe (A1×32-5 mm-25-177; 32-channel probe with 25-um spacing) into the brain, until the electrode would span the depths of ca. 800-1600 μm from the dura – thereby covering superficial SC. We waited about 15 minutes for the electrode to stabilize inside the brain before we started recording. The electrical signal from the electrodes was amplified and sampled at 24.4 kHz using a Tucker-Davis Technologies recording system.

First, we probed visual responses using a checkerboard stimulus consisting of black and white checkers of 20 visual degrees, that was displayed for 250 ms, then reversed for 250 ms, and was followed by a grey screen during the 1 s ITI. We then measured the RF of the recording sites using a sparse noise stimulus consisting of either 4 or 12 squares (50% black, 50% white) of 5 visual degrees at random locations on a grey background, that were displayed for 0.5 s followed by a 0.5 s ITI. This stimulus was shown for a total of 5-10 minutes. Using the receptive field data, we could ensure that the figure stimuli during the task were placed either inside or outside of the receptive field of the recorded sites. For the ‘figure’ stimulus, the figure was placed over the RF; for the ‘ground’ stimulus, the figure was placed 50-60 visual degrees lateral of the receptive field, in the hemifield contralateral to the RF (**Fig. 2F**). We proceeded to let the mouse perform the task while recording neuronal responses. After recording, we first disconnected the grounding and reference pins and shielding material close to the probe. We then removed the electrode from the brain and once again cleaned the craniotomy with sterile saline, and then sealed the craniotomy with Kwik-cast.

For our analysis of the electrophysiology data, we only included behavior sessions with good performance. Therefore, we tested whether the accuracy of the mouse on each variation of the task (i.e. contrast, orientation, phase) was significantly above chance level using a binomial test. If the session was shorter than 40 trials (the threshold for reaching statistical significance with 65% performance), we included the session if task accuracy was at least 65% and task accuracy on each side (i.e. figure stimulus on the left or right) was at least 50%.

To ensure the image was stable on the retina during the task, we excluded trials with eye movements. Given that the mice generally did not make many eye movements during the task, we excluded trials where the eye speed in the period between 0 and 450 ms after stimulus onset was higher than the mean speed + 2.5*SD.

For the identification of artefacts, we used an estimate of the envelop multi-unit activity (eMUA). The raw data was band-pass filtered between 500-5000 Hz, half-wave rectified (negative becomes positive) and then low-pass filtered at 200 Hz. The resulting signal constitutes the envelope of high-frequency activity. Each channel’s envelope signal was first z-scored across all trials *j* and samples *i* and the absolute value was taken to produce *zmua_ij_.* To identify samples at which the majority of recording-channels showed large excursions from the mean we took the geometric mean of *zmua_ij_* across all recording channels to produce Z_ij_:

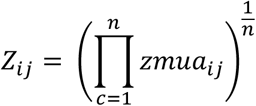

where *c* is the identity of the recording channel and *n* the total number of recording channels. Z_ij_ will be a positive number reflecting the consistency and extremeness of excursions from the mean across recording channels. As the geometric mean was used, Z_ij_ can only reach extreme values if the majority of recording channels show large excursions from the mean. We identified samples at which Z_ij_ was greater than 3 and removed these samples from all channels as well as removing the preceding and following 3 samples. We then recalculated Z_ij_ after removal of the extreme samples. To identify trials with extreme mean values (likely due to muscle artefacts) we took the mean value of Z_ij_ for each trial *j* and squared it to produce *χ_j_*:

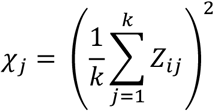

where *k* was the total number of trials. The distribution of *χ_j_* across trials was approximately normal and we fit the resulting distribution with a Gaussian function using non-linear least-squares fitting (using *fminsearch.m* in MATLAB) with mean μ and standard deviation σ. Extreme trials were identified as trials with *χ_j_* values more than 3σ from μ and were removed.

After applying the inclusion and exclusion criteria described above, we analyzed the single unit responses. First, we subtracted the common average across channels from the raw ephys data to reduce noise. The data was then further preprocessed and spike sorted using Kilosort (https://github.com/MouseLand/Kilosort; Pachitariu et al., 2016), with a spike detection threshold of −2 SD. The spike sorting results were manually curated using Phy (https://github.com/cortex-lab/phy). Spike clusters with high Kilosort quality scores that were stable across the session were included for analysis of single unit responses. We convolved the detected spikes of each unit with a Gaussian with an SD of 10 ms to derive a continuous estimate of the spike rate.

To estimate the receptive field of each neuron, we averaged the spikes that were evoked by each RF map checker in a time window in between 40 to 300 ms after checker onset. Given the variety of PSTH shapes, each neuron was assigned its own time window where the neuron showed increased or decreased spiking. We then fit a two-dimensional (2D)–Gaussian to estimate the width and center of both the ON and OFF RF. The quality of the fit was assessed using r^2^ and a bootstrapped variability index (BVI), which estimated the reliability of the RF center estimate (Kirchberger et al., 2021). We resampled an equal number of trials as in the original dataset (with replacement) and regenerated the Gaussian fit. The BVI is defined as the ratio of the SD of the RF center position and the SD of the fitted Gaussian. We used the most reliable fit of the RF (ON or OFF) as our RF estimate.

To investigate visually responsive neurons (**Fig. 2-4**), we included cells with an evoked response of at least 3 spikes/s in the period from 50 to 200 ms after stimulus onset. For investigating putative multisensory neurons (**Fig. 5** and **S3**), we included cells that had their peak firing rate between 650-900 ms after stimulus onset and had an task-evoked firing rate larger than 1 SD of their baseline response. Responses of each individual neuron were normalized to the mean response of that neuron across all trials where a grating was displayed inside the RF: *R_normalized_* = (*R* – *R_baseline_*)/(*R_max_* – *R_baseline_*), where *R* is the rate. The mean responses across neurons were smoothed by either 20 ms (figures displaying the visual response only) or 80 ms (figures displaying the entire trial).

We tested the difference between figure and ground (**Fig. 2I** and **3B**) based on an approach by Maris & Oostenveld (2007). In brief, the data is bootstrapped to make surrogate data-sets with randomly swapped condition labels. For each surrogate, we cluster together time points with significant p-values from a paired t-test and then take the maximum summed t-statistic across all clusters as a statistic. This builds up a null distribution of maximum cluster t-statistics. We then compare the cluster t-statistics from the unshuffled data to identify significant clusters. We estimated the latency of the figure-ground modulation by fitting a function (Poort et al., 2012) to the figure minus background response in a time window from 0 to 300 ms after stimulus onset. Briefly, the function is the sum of an exponentially modulated Gaussian and a cumulative Gaussian, capturing the Gaussian onset of neural modulation across trials/neurons and the dissipation of modulation over time. The latency was defined as the (arbitrary) point in time at which the fitted function reached 33% of its maximum value.

To further investigate the neural code in SC, we next tried to decode the trial identities from the neural responses we recorded. Generally, decoder algorithms need balanced data sets as input, to ensure non-biased training. However, our neural data was recorded during different behavioral sessions. Therefore we did not have balanced trial numbers across conditions for each neuron. Hence, we created a surrogate data set from our data to decode the stimulus type (figure vs. ground, **Fig 3**). For the surrogate data set of each task, we included only neurons for which we recorded at least 5 trials for each of the stimulus types (figure/ground). For each of the neurons, we excluded one random trial (either a figure trial of all neurons or a ground trial of all neurons). These trials together comprised the test set. We then pseudorandomly drew, with replacement, 10 trials of each stimulus type from each neuron’s remaining data. These comprised the training set. We then used a script derived from MATLAB’s Classification learner app to generate a linear SVM model that predicts the stimulus type based on the training data, and subsequently used that model to decode the test set. We repeated this 2000 times, with balanced test sets, for each time window; the performance was computed using a sliding window of 50 ms in steps of 10 ms. The resulting mean performances are reported. We tested whether the decoding performances were significantly different from chance using binomial tests with Bonferroni-Holm correction.

The d-prime was used to quantify the discriminability between figure and ground responses; it is a measure for the reliability of the signal on individual trials:

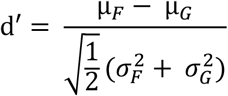

where μ*_F_* and μ*_G_* are the means, and 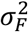 and 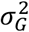 are the variances of the figure and ground response across trials, respectively. In **Figure 3**, we analyze d-prime values for the time window with the best decoding performance of each task. The shuffled data was generated by shuffling the trial identities of the real data 1000 times. In Figure 4 and 5, we analyze d-prime values (and firing rates) for the time window between the stimulus onset and the lick. To estimate the significance of the d-prime difference between hits and errors, we fit the data with a linear mixed-effects model. We selected the best model out of the possible models containing 1) three fixed effects (task, i.e. orientation or phase; response, i.e. hit or error; and intercept); 2) in- or excluding fixed effect interactions; and 3) in- or excluding random intercept terms to account for covariability of data obtained from the same unit, penetration and/or mouse. We defined the best model – balancing model fit and complexity - using the Akaike information criterion (AIC)(Akaike, 1974; Aho et al., 2014).

### Histology

We deeply anesthetized the mice with Nembutal and transcardially perfused them with phosphate-buffered saline (PBS), followed by 4% paraformaldehyde (PFA) in PBS. We extracted the brain and post-fixated it overnight in 4% PFA before moving it to a PBS solution. We cut the brains into 75-um-thick coronal slices and mounted them on glass slides in Vectashield DAPI solution (Vector Laboratories). We imaged the slices on either a Zeiss Axioplan 2 microscope (10× objective, Zeiss Plan-Apochromat, 0.16 NA) using custom-written Image-Pro Plus software or a Zeiss Axioscan.Z1 using ZEN software. The resulting histology images were used to confirm the location of fiber implants, the electrode trace and/or virus expression.

For estimating the histological depth of the neurons that we recorded during the electrophysiology experiments, we used a combination of electrophysiological and histological data. After outlier removal (see above) and common average subtraction of the raw data we low-pass filtered the data to get the local field potential (LFP). 50 Hz artifacts were removed by digital notch filtering. We computed the current source densities (CSD) from the LFP as described in Self et al. (2014). The CSD of the sSC typically showed one strong sink during the peak of the visual response. We therefore took the channel that had recorded the lowest value of the CSD as a reference for the relative position of the recording electrode in the sSC. To get an estimate of the absolute depth from the sSC surface, we measured the depths of the electrode tracks in the histological slices using ImageJ. From this, we estimated that the channel with the strongest CSD sink was located ca. 119 ± 48 μm from the SC surface. This value was combined with the information from the CSD to compute our estimate of the absolute depths of the recorded neurons. We tested the difference between the two groups using one-sample F test (for the difference between variances) and a Mann-Whitney U-test (for the difference between means)

## Acknowledgements

We thank Christiaan Levelt for sharing experimental facilities and Emma Ruimschotel for genotyping. We further thank Enny van Beest, Mehran Ahmadlou, Chris van der Togt and Ulf Schnabel for sharing their expertise. J.L.C and J.A.H. were funded by FLAG-ERA grant CHAMPmouse through de Nederlandse Organisatie voor Wetenschappelijk Onderzoek (NWO).

## Competing interests

The authors declare no competing interests.

## Notes

### Competing Interest Statement

The authors have declared no competing interest.

