## Supplementary Materials for "Involvement of superior colliculus in complex figure detection of mice"

### Supplementary analysis

#### Anesthetized electrophysiology for assessing strength of optogenetic inhibition

Mice were anesthetized by an intraperitoneal (IP) injection of 1.2 g urethane per kg body weight, supplemented by an IP injection of 8 mg chlorprothixene per kg body weight. We injected atropine sulfate (0.1 mg per kg, s.c.) to reduce mucous secretions. Additional doses of urethane were injected when a response to a toe-pinch was observed. Temperature was measured with a temperature probe underneath the mouse and maintained by a feedback-controlled heating pad set to 36.5 degrees. We cleaned the throat so the mouse could breathe freely and then head fixed the mouse by ear and bite bars. We applied lidocaine on the skin and then made an incision. After applying lidocaine on the periosteum, we drilled a 1 mm craniotomy for the electrode and optic fiber at 0.5 mm anterior and 0.5 mm lateral from the lambda cranial landmark. For the reference site, we drilled a 0.5 mm craniotomy in the contralateral frontal bone plate. We first inserted the reference wire into the reference site. Then we inserted a 200 µm diameter optic fiber, ca. 400 µm lateral to the desired electrode location. The optic fiber was inserted at a 15 degree angle to a depth of ca. 750 µm. We recorded using a laminar silicon electrode (A1 × 16-5mm-50-177-A16, 16 channels spaced 50 µm apart, Neuronexus). The electrode was coated with a green neuro-DiO (Biotium) or a red diI (Fisher Scientific) tracer solution and then inserted into the brain to a depth of ca. 1600 µm. We used black tin foil (Thorlabs) for shielding; this reduced external noise on the electrode as well as prevented laser light from reaching the mouse's eyes. We waited at least 15 minutes for the brain to stabilize before starting the recording. The electrical signal from the electrodes was amplified and sampled at 24.4 kHz using a Tucker-Davis Technologies recording system.

In order to assess the receptive field location of the recorded sites, we presented a 5 min movie (5 frames per second) of small black squares (approximately 5 degrees wide) in random positions on a white background (ratio of black to white area: 1:14 to 1:34) (Heimel et al., 2010). Then, to assess the effect of optogenetic inhibition, we displayed square wave gratings drifting in 8 different directions, with a spatial frequency of 0.02-0.05 cpd and a temporal frequency of 2 Hz. ITI duration was 1 second. Trials with and without optogenetic stimulation were interleaved. For optogenetic stimulation during stimulus presentation, we used either a 473 nm BL473T8-100FC laser or a 462 nm BLM462TA-100F laser (Shanghai Laser & Optics Century Co.), with a power of 4 mW at the fiber tip.

Analysis was done using custom Matlab scripts. The recorded signals were band-pass filtered (500-5000 Hz) and thresholded at 2x standard deviation to isolate spikes. All spikes from the same site were pooled together to get multi-unit activity. This was done using a fork from code written by Steve Van Hooser available online at https://github.com/heimel/InVivoTools. The spontaneous rate was defined as the mean rate in the last 0.5 s before stimulus onset. The evoked visual response was the mean rate during 0-0.5 s after stimulus onset, minus the spontaneous rate. The minimum evoked visual response for a unit to be included was 2 Hz. We used a paired t-test to assess the significance of the difference between the laser ON and laser OFF condition.

#### Eye movement analysis

To investigate whether the responses of the putative multisensory cells were correlated with eye movements, we z-scored both the neural responses and the eye movement speed. For the plots in **Fig. S3A**, we averaged the z-scored data across trials for one example neuron. For the analysis of the correlation in **Fig. S3B**, we concatenated the data across all the trials (i.e. creating one long time series) and computed the correlation coefficient between the eye movement and the neuronal responses. To test whether the resulting coefficients were significantly different from zero, we used a one-sample t-test ($p = 0.488$).

#### Supplementary Figures

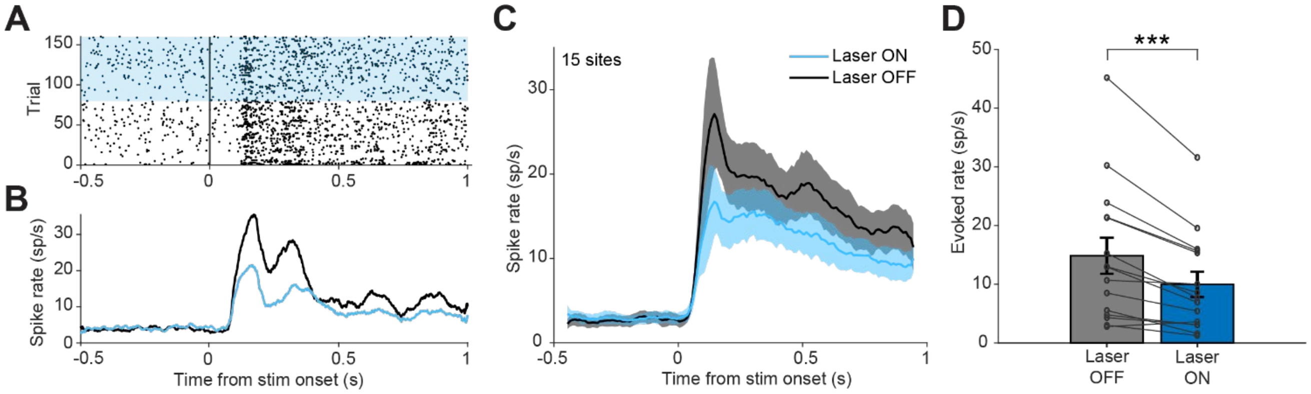


**Supplementary figure 1. Reduction of superior colliculus activity using optogenetic activation of GABA-ergic neurons. (A-B)** Responses of example neuron to drifting gratings (anesthetized electrophysiology). **(A)** Sorted raster plot. Each dot indicates a spike. Blue background indicates laser ON trials, white background laser OFF trials. Trials are sorted for presentation purposes. **(B)** Mean responses on laser ON (blue) and laser OFF (black) trials. **(C)** Population responses on laser ON (blue) and laser OFF (black) trials. Shading indicates SEM. **(D)** Visually evoked responses were significantly reduced in laser ON trials, by 33% on average. Bars indicate mean ± SEM. ***: p<0.001.

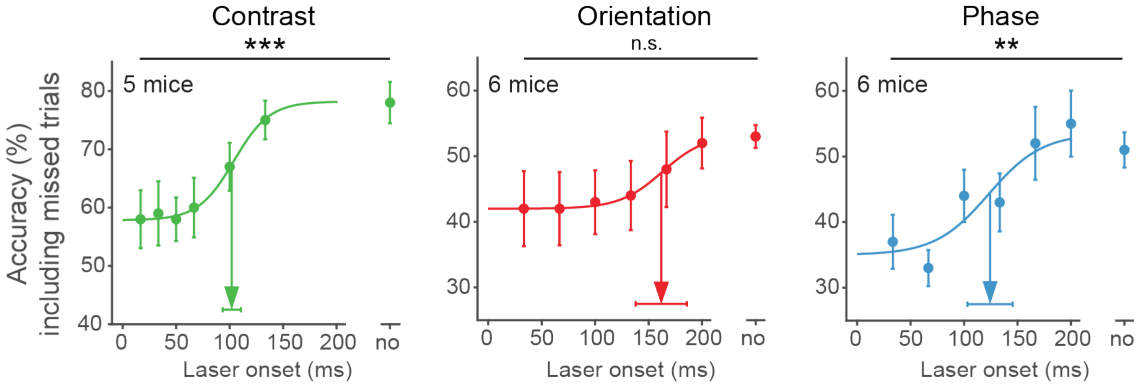


**Supplementary figure 2. Causal involvement of superior colliculus in figure detection tasks when including missed trials as error trials.** Inhibition of sSC significantly decreased task performance for the contrast and phase task. The accuracy on unperturbed trials without the laser condition is indicated by 'no'. Dots represent means ± SEM of accuracies across mice. Arrow and error bar indicate mean ± SD of bootstrapped fitted inflection points. Dashed line indicates chance level performance. *: p < 0.05, **: p < 0.01, ***: p<0.001.

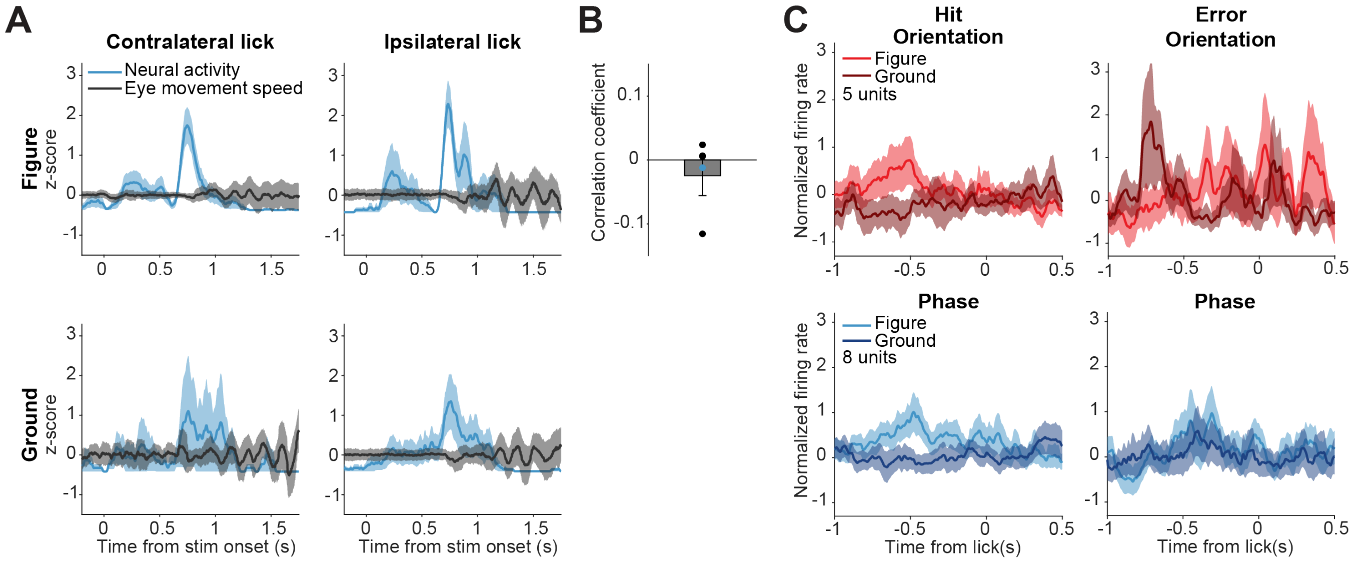


**Supplementary figure 3. Activity of putative multisensory neurons is not directly related to licking or eye movement.** **(A)** Example putative multisensory neuron. Mean neural activity plotted together with mean eye movement speed for different (stimulus/response) types of trials. Shading indicates SEM. **(B)** Correlation coefficients between eye movement speed and neural activity of putative multisensory neurons. Dots indicate individual neurons. Blue dot indicates the example neuron. **(C)** Population responses for hits vs. errors for all putative multisensory cells, centered on the time of the first lick in the response window. Note that the distinctive peak from **Fig. 5C** is not present here, suggesting that the peak response is related to the stimulus onset, and not to the first lick.

**Supplementary Table 1: Statistics**

**Figure 1**

| Figure panel | Comparison | Mean & SEM | Test | Tails or post-hoc | Test Statistic | p-value | Correction |
|---|---|---|---|---|---|---|---|
| 1E | Effect of optogenetics on contrast task (N = 5 mice) | 17 ms: 75% ± 3%<br>33 ms: 76 % ± 4%<br>50 ms: 74 % ± 4%<br>66 ms: 77% ± 6%<br>100 ms: 82 % ± 4%<br>133 ms: 91% ± 3%<br>No opto: 89 % ± 3% | One-way rm ANOVA | Post-hoc Tukey-Kramer | Main effect: F(6) = 5.16 | Main: p = 0.005 **<br>17 vs. no p = 0.027 *<br>33 vs. no p = 0.062<br>50 vs. no p = 0.022 *<br>66 vs. no p = 0.095<br>100 vs. no p = 0.551<br>133 vs. no p = 0.999 | Main: Bonferroni for 3 tasks |
| | Estimation inflection point - Contrast | 99 ± 8 ms (SD) | Bootstrap (n = 1000) | n.a. | | | |
| | Effect of optogenetics on orientation task (N = 6 mice) | 33 ms: 61% ± 2%<br>66 ms: 62 % ± 3%<br>100 ms: 60 % ± 3%<br>133 ms: 65% ± 4%<br>166 ms: 67% ± 3%<br>200 ms: 70% ± 3%<br>No opto: 73% ± 2% | One-way rm ANOVA | Post-hoc Tukey-Kramer | Main effect: F(6) = 3.55 | Main: p = 0.027 *<br>33 vs. no p = 0.034 *<br>66 vs. no p = 0.051<br>100 vs. no p = 0.021 *<br>133 vs. no p = 0.322<br>166 vs. no p = 0.566<br>200 vs. no p = 0.975 | Main: Bonferroni for 3 tasks |
| | Estimation inflection point - Orientation | 156 ± 35 ms (SD) | Bootstrap | n.a. | | | |
| | Effect of optogenetics on phase task (N= 6 mice) | 33 ms: 60% ± 4%<br>66 ms: 52 % ± 4%<br>100 ms: 64 % ± 2%<br>133 ms: 63% ± 6%<br>166 ms: 69% ± 3%<br>200 ms: 71% ± 3%<br>No opto: 73% ± 2% | One-way rm ANOVA | Post-hoc Tukey-Kramer | Main effect: F(6) = 5.03 | Main: p = 0.003**<br>33 vs. no p = 0.115<br>66 vs. no p = 0.001 **<br>100 vs. no p = 0.435<br>133 vs. no p = 0.702<br>166 vs. no p = 0.976<br>200 vs. no p = 0.999 | Main: Bonferroni for 3 tasks |
| | Estimation inflection point – Phase | 134 ± 30 ms (SD) | Bootstrap | n.a. | | | |

**Figure 2**

| Figure panel | Comparison | Mean & SEM | Test | Tails or post-hoc | Test Statistic | p-value | Correction |
|---|---|---|---|---|---|---|---|
| 2I | Figure vs. Ground – Contrast (22 units from 3 mice) | n.a. (too many comparisons) | Clustered T-statistic | One-tailed | T(22) | P<0.05 from 68-99 ms | Clustered T-statistic, see Methods section |
| | Figure vs. Ground – Orientation (49 units from 5 mice) | n.a. (too many comparisons) | Clustered T-statistic | One-tailed | T(48) | P<0.05 from 84-129 ms | Clustered T-statistic, see Methods section |
| | Figure vs. Ground – Phase (32 units from 5 mice) | n.a. (too many comparisons) | Clustered T-statistic | One-tailed | T(31) | No significant clusters | Clustered T-statistic, see Methods section |
| 2J | Onset modulation - Contrast | 67 ms | Curve fitting | n.a. | | | |
| | Onset modulation - Orientation | 75 ms | Curve fitting | n.a. | | | |

**Figure 3**

| Figure panel | Comparison | Mean & SEM | Test | Tails or post-hoc | Test Statistic | p-value | Correction |
|---|---|---|---|---|---|---|---|
| 3B | Figure vs. Ground – Orientation (12 units from 3 mice) | n.a. (too many comparisons) | Clustered T-statistic | One-tailed | T(11) | P<0.05 from 81-102 ms | Clustered T-statistic, see Methods section |
| | Figure vs. Ground – Phase (14 units from 3 mice) | n.a. (too many comparisons) | Clustered T-statistic | One-tailed | T(13) | No significant clusters | Clustered T-statistic, see Methods section |
| 3C | Orientation decoder | n.a. (too many comparisons) | Binomial test | Two-tailed | n.a. (N = 2000) | 0-50 ms: p = 1.519<br>10-60 ms: p = 0.468<br>20-70 ms: p = 1.257<br>30-80 ms: p = 1.230<br>40-190 ms: p <0.001 ***<br>150-200 ms: p = 0.024 *<br>160-210 ms: p = 1.302<br>170-220 ms: p = 1.519<br>180-230 ms: p = 1.006<br>190-240 ms: p = 0.025 *<br>200-250 ms: p = 0.230<br>210-260 ms: p = 0.002 **<br>220-270 ms: p = 0.771<br>230-280 ms: p = 1.033 | Bonferroni-Holm |

| | | | | | | | |
|---|---|---|---|---|---|---|---|
| | Phase decoder | n.a. (too many comparisons) | Binomial test | Two-tailed | n.a.<br>(N = 2000) | 0-50 ms: p = 0.345<br>10-60 ms: p = 0.118<br>20-70 ms: p < 0.001 ***<br>30-80 ms: p = 0.336<br>40-90 ms: p < 0.001 ***<br>50-100 ms: p < 0.001 ***<br>60-110 ms: p = 0.126<br>70-120 ms: p = 0.345<br>80-130 ms: p = 0.257<br>90-140 ms: p = 0.336<br>100-150 ms and later bins: p < 0.001 *** | Bonferroni-Holm |
| 3D | Relative model weight of RF inside vs. RF edge Neurons – Orientation<br>(49/23 units from 5/3 mice) | RF inside:<br>0.214 ± 0.035<br>RF edge:<br>0.220 ± 0.064 | Mann-Whitney U-test | Two-tailed | Z = 0.172<br>Ranksum = 382 | P = 0.863 | None (all n.s.) |
| | Relative model weight of RF inside vs. RF edge Neurons – Phase<br>(32/14 units from 5/3 mice) | RF inside:<br>0.049 ± 0.059<br>RF edge:<br>0.149 ± 0.069 | Mann-Whitney U-test | Two-tailed | Z = 1.086<br>Ranksum = 375 | P = 0.277 | None (all n.s.) |
| 3E | Orientation mean d’ | Mean d’ = 0.302 | Bootstrap (n = 5000) | Two-tailed | CI 99.9%<br>-0.147 – 0.137 | P < 0.001 *** | None |
| | Phase mean d’ | Mean d’ = 0.090 | Bootstrap (n = 5000) | Two-tailed | CI 95%<br>-0.085 – 0.085 | P < 0.05 * | None |

**Figure 4**

| Figure panel | Comparison | Mean & SEM | Test | Tails or post-hoc | Test Statistic | p-value | Correction |
|---|---|---|---|---|---|---|---|
| 4C | d-prime for<br>- Hit vs. Error<br>- Orientation vs. Phase<br><br>(61/54 and 46/46 units for orientation hit/error and phase hit/error, respectively) | Orientation Hit:<br>0.404 ± 0.078<br>Orientation Error:<br>-0.093 ± 0.138<br>Phase Hit:<br>0.325 ± 0.098<br>Phase Error:<br>0.158 ± 0.131 | LME:<br>*dprime ~ 1 + Respone + Task + Response * Task + (1\|Unit)* | Matlab LME standard | **Main:**<br>Response: F(1,190) = 12.089<br>Task: F(1,190) = 0.303<br>Response*Task: F(1,190) = 2.589<br><br>**Post Hoc:**<br>Response (Orientation)<br>T(101) = 3.537<br>Response (Phase)<br>T(89) = 1.036 | **Main:**<br>P <0.001 ***<br>P = 0.583<br>P = 0.109<br><br>**Post-Hoc :**<br><br>P < 0.001 ***<br><br>P = 0.303 | None |

**Figure 5**

| Figure panel | Comparison | Mean & SEM | Test | Tails or post-hoc | Test Statistic | p-value | Correction |
|---|---|---|---|---|---|---|---|
| 5D | d-prime for<br>- Hit vs. Error<br>- Orientation vs. Phase<br><br>(5/5 and 8/8 units for orientation hit/error and phase hit/error, respectively) | Orientation Hit:<br>0.671 ± 0.429<br>Orientation Error:<br>-0.353 ± 0.496<br>Phase Hit:<br>0.607 ± 0.439<br>Phase Error:<br>-0.398 ± 0.491 | LME:<br>*dprime ~ 1 + Response + Task* | MATLAB LME standard | **Main:**<br>Response: F(1,19) = 4.645<br>Task: F(1,19) = 0.014<br><br>**Post Hoc:**<br>Response (Orientation)<br>T(7) = 1.569<br>Response (Phase)<br>T(111) = 1.480 | **Main :**<br>P = 0.044*<br>P = 0.906<br><br>**Post-Hoc :**<br><br>P = 0.161<br><br>P = 0.167 | None |
| 5E | Variance of depth of visual vs. MS cells<br><br>Difference in depth of visual vs. MS cells<br><br>(99 and 8 units for visual and MS cells, respectively) | Visual: 338 ± 190 um (SD)<br>Multisensory: 416 ± 92 um (SD) | Two-sample F-test<br><br>Mann-Whitney U-test | Two-tailed<br><br>Two-tailed | F(7, 98) = 0.231<br><br>Z = 1.352 | P = 0.046<br><br>P = 0.177 | None<br><br>None |

**Figure S1**

| Figure panel | Comparison | Mean & SEM | Test | Tails or post-hoc | Test Statistic | p-value | Correction |
|---|---|---|---|---|---|---|---|
| S1D | No-opto vs. Opto<br>(15 sites from 3 mice) | No-opto: 14.8 ± 3.1 sp/s<br>Opto: 10.0 ± 2.2 sp/s | Paired t-test | Two-tailed | T(14) = 4.81 | P < 0.001 *** | none |

**Figure S2**

| Figure panel | Comparison | Mean & SEM | Test | Tails or post-hoc | Test Statistic | p-value | Correction |
|---|---|---|---|---|---|---|---|
| S2 | Effect of optogenetics on contrast task – including missed trials<br>(N = 5 mice) | 17 ms: 58% ± 4%<br>33 ms: 59% ± 5%<br>50 ms: 58 % ± 3%<br>66 ms: 60% ± 5%<br>100 ms: 67 % ± 4%<br>133 ms: 75% ± 3%<br>No opto: 78 % ± 3% | One-way rm ANOVA | Post-hoc Tukey-Kramer | Main effect:<br>F(6) = 9.10 | Main: p < 0.001 ***<br>17 vs. no p < 0.001 ***<br>33 vs. no p < 0.001 ***<br>50 vs. no p < 0.001 ***<br>66 vs. no p = 0.002 **<br>100 vs. no p = 0.115<br>133 vs. no p = 0.972 | Main:<br>Bonferroni for 3 tasks |

| | | | | | | | |
|---|---|---|---|---|---|---|---|
| | Estimation inflection point including missed trials - Contrast | 102 ± 8 ms (SD) | Bootstrap (n = 1000) | n.a. | | | |
| | Effect of optogenetics on orientation task - including missed trials<br>(N = 6 mice) | 33 ms: 42% ± 5%<br>66 ms: 42% ± 5%<br>100 ms: 43 % ± 4%<br>133 ms: 44% ± 5%<br>166 ms: 48% ± 5%<br>200 ms: 52% ± 4%<br>No opto: 53% ± 2% | One-way rm ANOVA | Post-hoc Tukey-Kramer | Main effect: F(6) = 2.12 | Main: p = 0.24<br>33 vs. no p = 0.253<br>66 vs. no p = 0.213<br>100 vs. no p = 0.342<br>133 vs. no p = 0.461<br>166 vs. no p = 0.912<br>200 vs. no p = 1.000 | Main: Bonferroni for 3 tasks |
| | Estimation inflection point including missed trials - Orientation | 161 ± 25 ms (SD) | Bootstrap | n.a. | | | |
| | Effect of optogenetics on phase task – including missed trials<br>(N = 6 mice) | 33 ms: 37% ± 4%<br>66 ms: 33% ± 3%<br>100 ms: 44% ± 4%<br>133 ms: 43% ± 4%<br>166 ms: 52% ± 5%<br>200 ms: 55% ± 5%<br>No opto: 51% ± 2% | One-way rm ANOVA | Post-hoc Tukey-Kramer | Main effect: F(6) = 5.6 | Main: p = 0.002**<br>33 vs. no p = 0.099<br>66 vs. no p = 0.016 *<br>100 vs. no p = 0.829<br>133 vs. no p = 0.651<br>166 vs. no p = 1.000<br>200 vs. no p = 0.973 | Main: Bonferroni for 3 tasks |
| | Estimation inflection point including missed trials – Phase | 123 ± 21 ms (SD) | Bootstrap | n.a. | | | |

**Figure S3**

| Figure panel | Comparison | Mean & SEM | Test | Tails or post-hoc | Test Statistic | p-value | Correction |
|---|---|---|---|---|---|---|---|
| S3B | Correlation of eye speed with neural activity vs. correlation coefficient of zero<br>(N = 4 units) | Coefficients: -0.025 ± 0.031 | One-sample t-test | Two-tailed | T(3) = -0.789 | P = 0.488 | None |
